# A Novel Blood-Based Epigenetic Clock for Intrinsic Capacity Predicts Mortality and is Associated with Clinical, Immunological and Lifestyle Factors

**DOI:** 10.1101/2024.08.09.607252

**Authors:** Matías Fuentealba, Laure Rouch, Sophie Guyonnet, Jean-Marc Lemaitre, Philipe de Souto Barreto, Bruno Vellas, Sandrine Andrieu, David Furman

## Abstract

Age-related decline in intrinsic capacity (IC), defined as the sum of an individual’s physical and mental capacities, is a cornerstone for promoting healthy aging and longevity, as it emphasizes maximizing function throughout the aging process instead of merely treating diseases. However, accurate assessments of IC are resource-intensive, and the molecular and cellular basis of its decline are poorly understood. Herein, we used the INSPIRE-T cohort, consisting of 1,014 individuals aged 20 to 102, to construct the IC clock, a DNA methylation (DNAm)-based predictor of IC trained on the clinical evaluation of cognition, locomotion, psychological well-being, sensory abilities, and vitality. In the Framingham Heart Study, age-adjusted DNAm IC correlates with first- and second-generation epigenetic clocks, predicts all-cause mortality, and is strongly associated with changes in molecular and cellular immune and inflammatory biomarkers, functional and clinical endpoints, health risk factors, and diet.

## Main

In 2015, the World Health Organization (WHO) introduced the concept of intrinsic capacity (IC), defined as the sum of all physical and mental capacities that an individual can draw on at any point in their life^1^. This concept serves as a foundation for assessing and promoting healthy aging and longevity, with the goal of shifting healthcare’s focus from identifying and treating acute episodes of illness toward quantifying and maintaining functional ability as people age^1–3^. Although IC varies greatly between individuals, it typically peaks in early adulthood and declines after midlife, and adopting a healthier lifestyle (such as physical exercise and good nutrition) can improve IC at any age^4–9^.

The International Classification of Diseases (11th Revision) (ICD-11) recently added the ‘age-related decline in IC’ under code MG2A^10^, allowing practitioners to use IC as a standard metric of functional aging and encouraging the development of personalized therapies that promote healthy aging by maintaining optimal IC levels. Since the inception of IC, dozens of studies have developed IC scores and validated its association with various health-related factors (recently reviewed by Beyene et al.^11^). For example, in our latest study on age-specific reference centiles for IC using the INSPIRE (Institute for Prevention Healthy Aging and Medicine Rejuvenative) human lifespan translational (T) cohort (INSPIRE-T cohort), we showed that individuals with low IC had significantly higher comorbidity, frailty, difficulties in activities of daily living, and more falls^12^.

Despite the advantages of using IC to assess functional ability as we age, (1) current methods to quantify it require various equipment and trained personnel to administer the tests accurately, and (2) we still do not understand the molecular and cellular mechanisms underlying the age-associated decline in IC. To address these, we collected DNA methylation (DNAm) data from participants in the INSPIRE-T cohort to construct an epigenetic predictor of IC (IC clock). Then we applied the IC clock to the Framingham Heart Study (N = 2503) and evaluated the association between changes in DNAm IC (age-adjusted) and mortality, clinical markers of health and lifestyle. We also establish the molecular mechanisms of IC by determining the gene expression and cell composition changes associated with changes in DNAm IC.

## Results

### Intrinsic capacity declines with age

Using clinical assessment tools, we developed an IC score that represents the combined age-related decline in five health domains: cognition, locomotion, sensory (vision and audition), psychological, and vitality (see Methods) (Fig. 1a). The IC scores ranged from 0 to 1, with 1 indicating the best possible health outcome and 0 indicating the worst. We examined the correlation between chronological age and each domain, as well as overall IC. We found a negative relationship between age and each domain of IC, with the strongest correlation being between age and the overall IC (r_s_ = -0.65, p = 9.97e-117) (Fig. 1b). The psychological domain showed the smallest correlation with age (r_s_ = -0.07, p = 1.94e-02).

**Figure 1.**
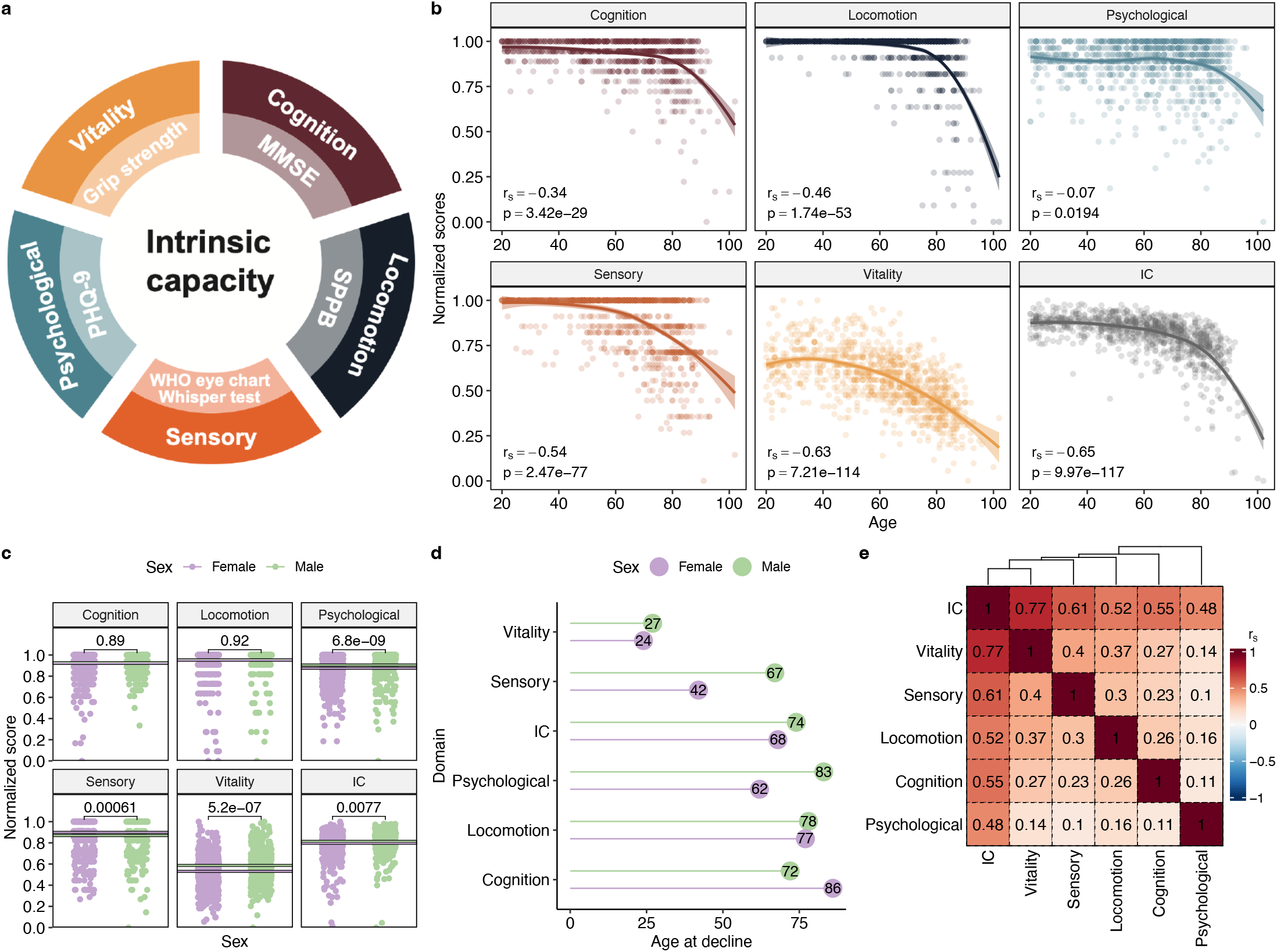
Age- and sex-related changes in intrinsic capacity. **a**, Domains of IC and clinical assessment tools used for the derivation of the IC score. **b**, Correlation between the scores in each domain (and overall) and chronological age. Due to its nonlinearity, correlation values were calculated using Spearman’s method. We estimated the regression line using the Locally Estimated Scatterplot Smoothing (LOESS) method. **c**, Average scores for each IC domain in males (green) and females (purple). Lines indicate the mean value for each sex in each domain. We calculated the p-values for the mean differences using a Wilcoxon’s test. **d**, Estimated age at decline obtained from a continuous two-phase model regression analysis in each domain. **e**, Correlation between each IC domain’s score. MMSE: Mini-Mental State Examination; SPPB: Short Physical Performance Battery; PHQ-9: Patient Health Questionnaire (for depression disorder).

Given the well-established differences in healthspan and lifespan between males and females, we investigated the sex differences in the severity and rates (age at decline) of IC domains. We found that males have higher average scores in the psychological and vitality domains (p = 6.80e-09 and 5.20e-07, respectively), while females have slightly higher scores in the sensory domain (p = 6.10e-04) (Fig. 1c). Cognition and locomotion showed no statistically significant differences between sexes. We used a continuous two-phase model regression analysis to estimate the average age at decline in each domain (see Methods). We found that, despite having higher sensory function on average, females exhibit an earlier decline (42 years old versus 67 years old in males) (Fig. 1d). In contrast, cognitive decline occurred earlier in males (72 years old) than in females (86 years old), suggesting that females tend to maintain cognitive resilience for longer.

We next assessed the contribution of each IC domain to the overall IC. To do so, we calculated the correlation between the scores for each domain and the overall IC score. We observed that the overall IC score had a stronger positive correlation with each domain (r_s_ between 0.48 and 0.77) than the correlations between domains, confirming the IC score’s integrative nature (Fig. 1e). The sensory and vitality domains had the highest inter-domain correlation (r_s_ = 0.4), while the psychological and sensory domains had the lowest (r_s_ = 0.1).

### DNA-methylation based predictor of intrinsic capacity

We used DNA methylation levels (Infinium EPIC array) from 933 INSPIRE-T participants to predict IC. To ensure cross-platform usability, we used data from 361,080 CpGs that passed quality control tests and overlapped with the 450K array (see Methods). We built the predictive model using elastic-net regression and 10-fold cross-validation^13^. We ranked models with different elastic net mixing parameters (alpha) based on the correlation between real and predicted values, the model’s error, and the number of CpGs used (Fig. 2a). The best model was the one with the best ranking across all three metrics (alpha = 0.9, 91 CpGs, r_s_ = 0.61, Mean Absolute Error (MAE) = 0.053) (Fig. 2b). Interestingly, DNAm-estimated IC had a stronger correlation with age (r_s_ = -0.92) than with the calculated IC score (Fig. 2c).

**Figure 2.**
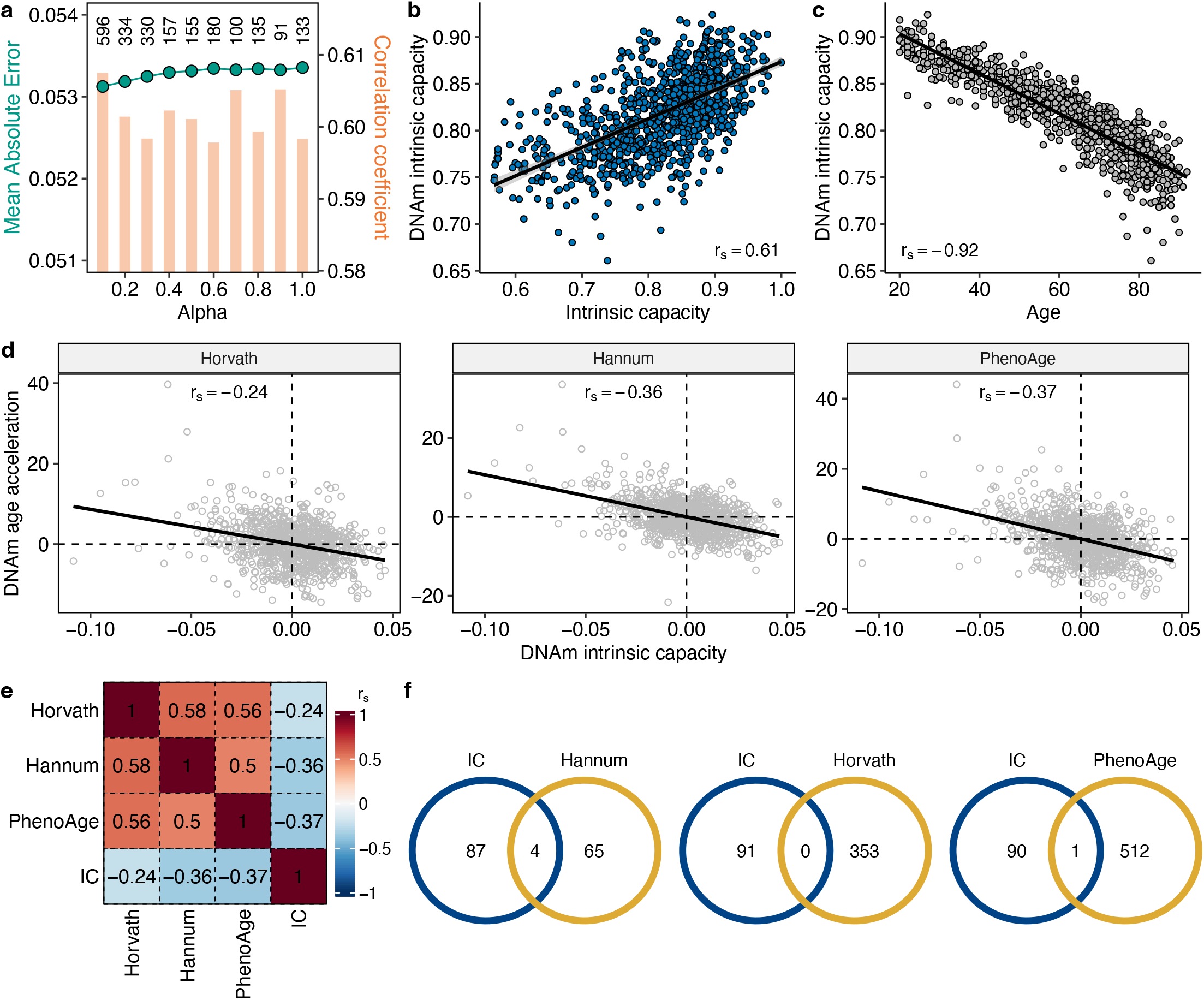
Model selection and correlation with other DNA methylation clocks. **a**, Performance of the model at values of alpha between 0.1 and 1. **b**, Correlation between IC and the DNA methylation-based estimate of IC (DNAm IC) in the best model. **c**, Correlation between DNAm IC and chronological age. **d**,**e** Correlations between age acceleration from epigenetic clocks (i.e. age-adjusted epigenetic age) and age-adjusted DNAm IC. **f**, Overlap between CpGs included in the IC clock (blue circles) and epigenetic clocks (yellow circles).

We compared the IC clock to first- and second-generation clocks designed to predict chronological age (Horvath and Hannum) or clinical markers and mortality (PhenoAge). As expected, we found a negative correlation between DNAm IC and epigenetic clocks (independent of chronological age), with PhenoAge showing the strongest correlation (r_s_ = -0.37, p < 0.001), followed by the Hannum clock (r_s_ = -0.36, p < 0.001) (Fig. 2d). Furthermore, the absolute magnitude of the correlation was higher within previous epigenetic clocks compared to DNAm IC (absolute mean r_s_ = 0.58 vs. 0.24) (Fig. 2e), suggesting that the IC clock captures a distinct aspect of aging biology, in line with the finding of no major overlaps between the CpG sites included in each clock and DNAm IC (Fig. 2f).

### DNAm intrinsic capacity is linked with immune response alterations

To better understand the molecular components and biological processes associated with DNAm IC, we calculated DNAm IC in the Framingham Heart Study (FHS). Using PBMC gene expression data available for 2,139 participants, we performed differential expression analysis with age-adjusted DNAm IC as the outcome. We found that the expression of 578 genes (286 up-regulated and 292 down-regulated) was significantly associated with changes in DNAm IC, independent of chronological age (Fig. 3a, Supplementary Table 1). Among the top genes, higher DNAm IC was strongly associated with a significant increase in expression of CD28 (False Discover Rate (FDR) = 1.06e-32), a surface molecule highly expressed in CD4^+^ and CD8^+^ T-cells whose loss of expression is a hallmark of immunosenescence. Additionally, we linked higher DNAm IC to increased expression of CAMK4, which controls follicular helper T-cell expansion^14^ and is essential for proper antibody production in response to vaccination in animal models and humans^15^. In contrast, poor IC clock levels tracked with elevated expression of PFTK1 (CDK14) (FDR = 2.76e-29), a regulator of the Wnt signal transduction pathway and proinflammatory mediator associated with Parkinson’s disease and a variety of cancers, and sensitive to changes in diet^16,17^ and Fibromodulin (FMOD) (FDR = 7.38e-27), a master regulator of inflammatory responses and inflammaging-related diseases such as poor wound healing, osteoarthritis, tendinopathy, atherosclerosis, and heart failure^18^. To further investigate these associations, we performed gene ontology enrichment analysis to identify associated biological processes. We found that most of the genes associated with age-adjusted DNAm IC are involved in the immune response, especially T-cell activation (Fig. 3b, Supplementary Table 2).

**Figure 3.**
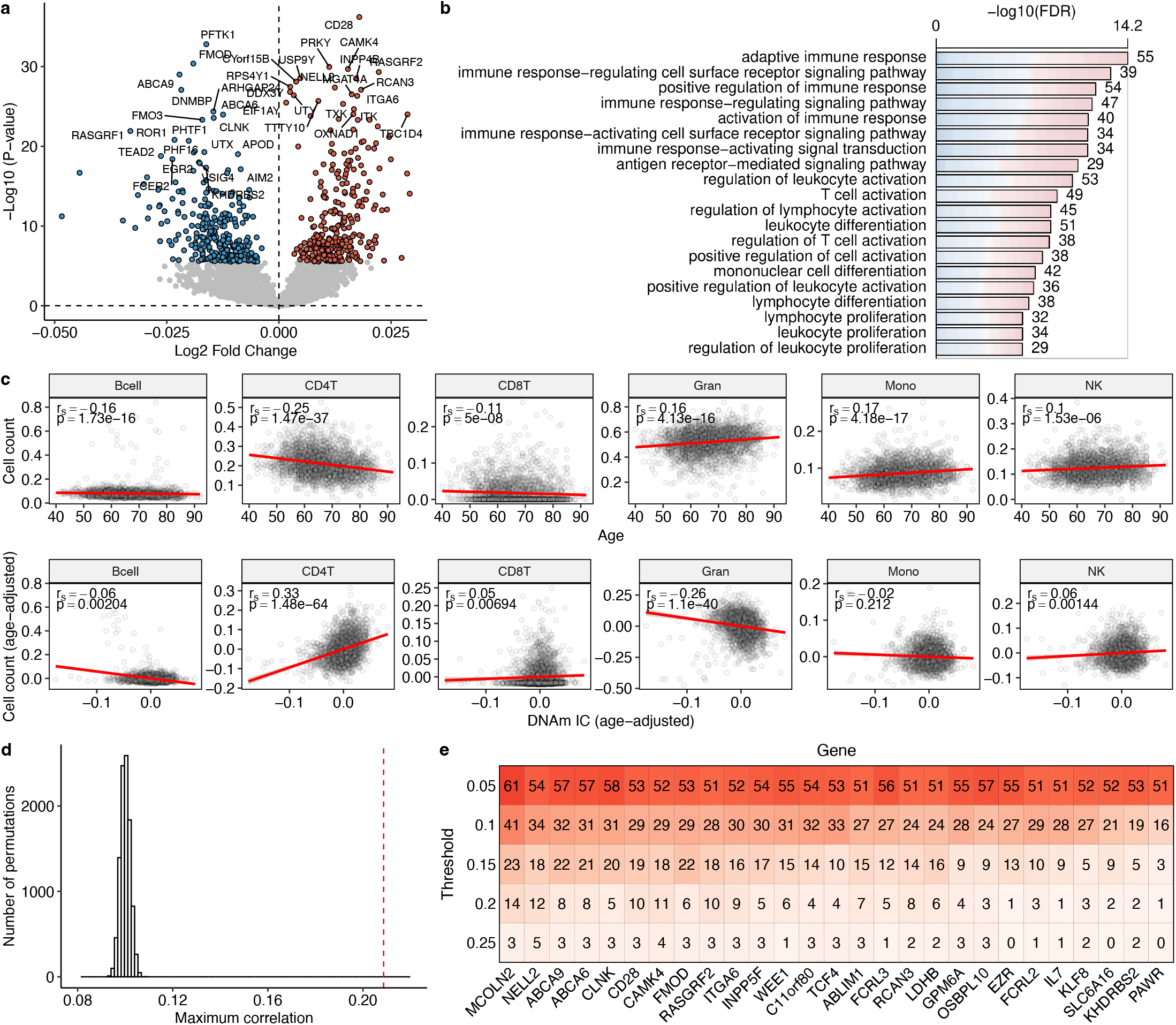
Age-Independent gene expression and immune cell frequency correlates of DNAm IC. **a**, Genes whose expression is significantly associated with age-independent changes in DNAm IC (colored dots). Labels indicate the top 10 genes in each direction. **b**, Top 20 biological processes enriched in genes significantly associated with DNAm IC. Cell frequency changes with age and DNAm IC. **c**, The top panels show the relationship between the DNAm-based cell count estimates and chronological age. The bottom panels display the relationship between DNAm IC and cell count independently of chronological age. Red lines indicate the linear regression between the values. **d**, Average maximum correlation between beta values from the IC clock CpGs and gene expression of genes associated with age-adjusted DNAm IC (dotted red line). The histogram indicates the values expected by chance based on 10,000 permutations of the gene set. **e**, Genes associated with more than 55% of the CpGs in the IC clock at a Pearson’s correlation threshold of 0.05 or higher. We ranked genes based on the average number of associated CpGs across all thresholds.

To better interpret these findings, we used the Houseman’s estimation method, which utilizes DNA methylation data to estimate blood cell counts, to examine the relationship between DNAm IC and cell populations (Fig. 3c). While the CD4^+^ T-cell frequency decreased significantly with age (r_s_ = -0.25, p = 1.47e-37), higher DNAm IC was associated with a greater number of CD4^+^ T-cells (r_s_ = 0.33, p = 1.48e-64). Given that CD28 is frequently found on the surface of CD4^+^ T-cells, this could explain why individuals with higher DNAm IC have higher levels of CD28 expression (see Discussion). Similarly, the number of granulocytes tended to increase with age (r_s_ = 0.16, p = 4.13e-16) and decrease in individuals with higher DNAm IC (r_s_ = -0.26, p = 1.1e-40).

Next, we analyzed the link between the genes with expression associated with DNAm IC and the CpGs in the IC clock by computing the correlation between the CpGs beta values (i.e. proportion of methylation ranging from 0 (unmethylated) to 1 (methylated)) and the gene expression values. We found that, on average, CpGs in the IC clock have a correlation of 0.21 with the expression of at least one significantly associated gene, whereas an average correlation of 0.1 is expected by chance (permutation p<0.0001) (Fig. 3d). This analysis confirms a strong connection between the methylation of the CpGs in the IC clock and the identified gene expression signature of DNAm IC.

We also examined the genes that showed absolute correlations above various thresholds and found that the expression of MCOLN2 correlates with beta values for 61 out of 91 CpGs (67%) (Fig. 3e, Supplementary Table 3). We observed that low expression of MCOLN2 was associated with higher DNAm IC. Interestingly, studies have found that MCOLN2 promotes virus entry and infection^19^ and activates the age-related NLRP3 inflammasome in response to oxidative stress, and contributes to the secretion of pro-inflammatory Interleukin-1 beta^20^. Also, CD28, the gene whose expression is most associated with age-adjusted DNAm IC, displayed a correlation with the methylation of more than half of the CpGs in the IC clock. Overall, the findings indicate that the IC clock detects aspects of immunosenescence in the blood that are associated with functional aging changes.

### DNAm IC is associated with all-cause mortality risk, clinical markers of health and lifestyle

Despite not developing the IC clock using mortality data, we hypothesized that the DNAm estimate of IC can also predict mortality, given the previous identification of IC as a mortality risk factor. Using FHS mortality data, we investigated whether DNAm IC was associated with an increased mortality risk from all causes or age-related conditions. We found that DNAm IC was slightly more significantly associated with all-cause mortality risk than PhenoAge (trained on mortality data), Horvath, and Hannum clocks (HR = 1.33, p = 1.05e-16) when comparing the quintiles with the highest (high IC) and lowest (low IC) age-adjusted DNAm estimates (Fig. 4a-b). In addition, DNAm IC was significantly associated with an increased risk of death from age-related diseases such as cardiovascular disease (HR = 1.23, p = 2.46e-05), congestive heart failure (HR = 1.26, p = 3.11e-04), and stroke/TIA (HR = 1.18, p = 2.55e-02). To help interpret these findings, we examined Kaplan-Meier survival curves for various causes of death. We estimated that a person with high DNAm IC lives on average 4.94 years longer than someone with low DNAm IC (Fig. 4c).

**Figure 4.**
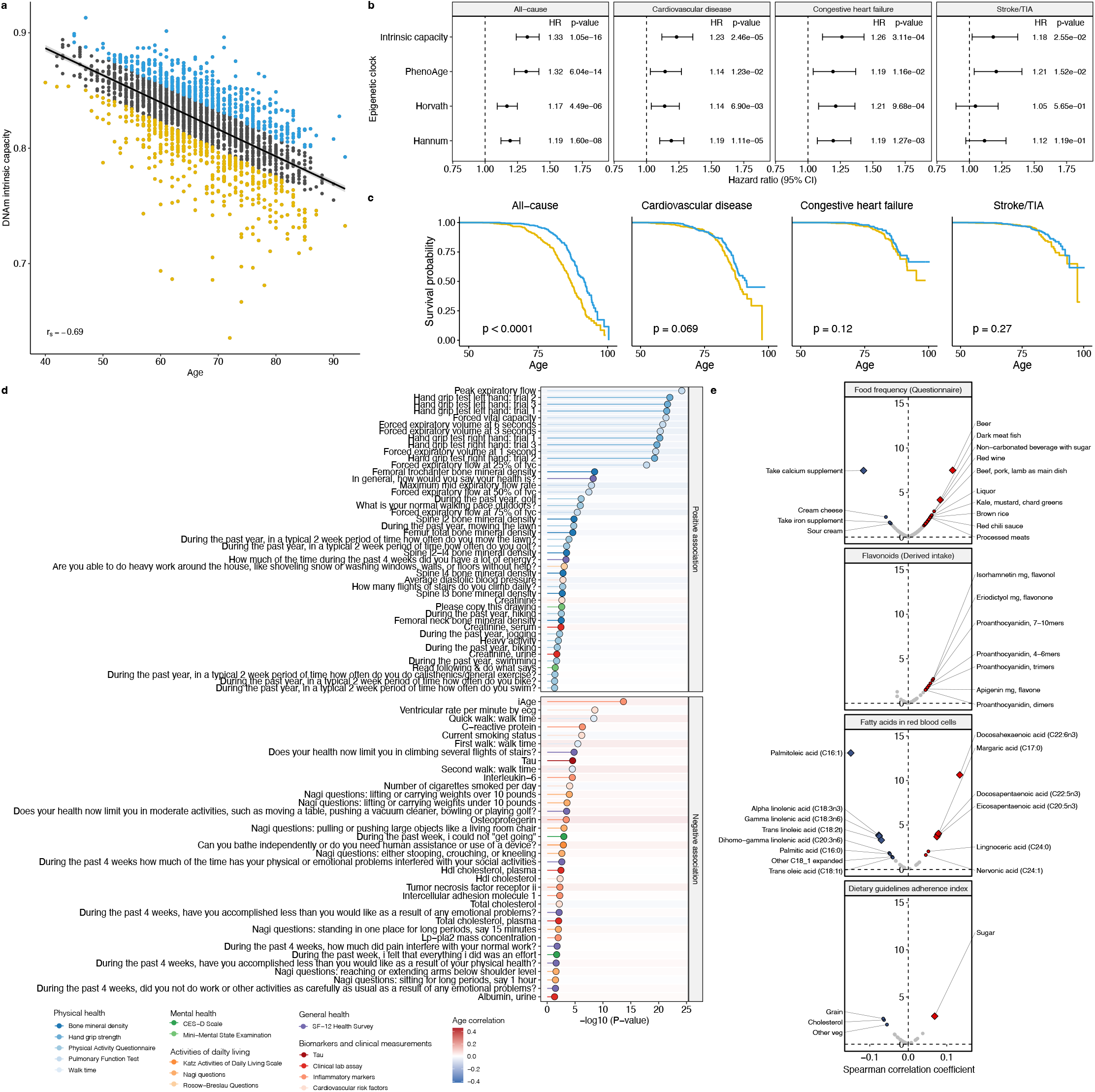
DNAm IC predicts mortality and is associated with functional, clinical, and lifestyle factors. **a**, Individuals with high (blue) and low (yellow) age-adjusted DNAm IC in the FHS. **b**, Forest plot shows the results from Cox proportional hazard models adjusting for chronological age for DNAm IC and first- and second-generation epigenetic clocks. **c**, Kaplan-Meier survival estimates for individuals with high and low DNAm IC (age-adjusted). P-values were calculated using a log-rank test. **d**, Significance of the association between health measurements and high or low DNAm IC using a logistic regression. Dot colors and lines indicate the data types in which the variable was measured. The background color for each variable indicates the correlation with age. **e**, Correlations between DNAm IC and consumption of different foods, derived intake of flavonoids, fatty acid concentrations in the blood, and dietary adherence. Rhombuses display significant correlations after multiple testing correction.

We also calculated the association between DNAm IC and assessments of physical and mental health, questionnaires of daily living activities, overall health and biomarkers, and clinical measurements. We found that individuals with high age-adjusted DNAm IC tend to display better pulmonary function (Fig. 4d, Supplementary Table 4). As expected, given its inclusion as a vitality indicator, we observed superior grip strength in individuals with high DNA IC. Also, faster walk time, greater bone mineral density, and better self-reported health perception were among the features most significantly associated with high DNA IC. We also observed a negative association between high DNAm IC and inflammatory age (iAge)^21^, canonical inflammatory markers such as CRP and IL-6, markers of neurodegeneration such as Tau, and smoking status.

Using the comprehensive food frequency questionnaire from the FHS, we explored the relationship between DNAm IC and specific food consumption. We found that individuals with higher DNAm IC tend to consume more beer (FDR = 4.48e-06) and dark meat fish (i.e. mackerel, salmon, sardines, bluefish, and swordfish) (FDR = 9.5e-03), but fewer calcium supplements (FDR = 4.84e-06) (Fig. 4e, Supplementary Table 5). We also analyzed the correlation with flavonoid intake derived from the food frequency questionnaire and found that the consumption of most flavonoids was associated with higher DNAm IC, but none specifically reached statistical significance after multiple testing correction. We also estimated whether there was an association with fatty acids in red blood cells measured by gas chromatography. We found that elevated concentrations of docosahexaenoic acid (FDR = 6.03e-10), docosapentaenoic acid (FDR = 2.95e-03), and eicosapentaenoic acid (FDR = 4.27e-03), all three long-chain omega-3 fatty acids from marine origin, were associated with higher DNAm IC. Lastly, we analyzed the dietary guideline adherence questionnaire and found that consuming sugar at the recommended level (<=5% of total energy) was associated with a higher DNAm IC (FDR = 3.01e-02). Overall, these results suggest that consuming fish rich in long-chain omega-3 fatty acids and adhering to recommended sugar intake guidelines is associated with IC maintenance.

## Discussion

In this study, we constructed a methylation-based clock to monitor age-related decline in IC, which predicts mortality, tracks cardiovascular risk factors and functional resilience, and is strongly associated with immune function and inflammatory health. While the association between IC and health-related factors is well-documented^11^, studies looking into the biological basis of IC are lacking. In this study, we revealed important relationships between DNAm IC and the immune system. As we age, both CD4^+^ and CD8^+^ T-cells gradually lose their ability to express CD28, a gene essential for T-cell activation and proliferation^22,23^, which ultimately results in immunosenescence and a reduced immune response in the elderly^24,25^. Notably, we observed that individuals with high DNAm IC display increased expression of CD28 and higher CD4^+^ T-cell population, suggesting that maintaining DNAm IC levels could be an effective strategy for preserving immune function with age.

We previously derived a metric for systemic chronic age-related inflammation (iAge) based on gene expression data from the 1000 Immunomes project (1KIP) cohort (n = 397)^21^ and showed that iAge was an important biomarker for immune and cardiovascular health. Herein, we found that individuals with high DNAm IC display significantly lower iAge. Consistently, we also observed negative associations between DNAm IC and the levels of canonical inflammatory markers such as CRP and IL-6, which have also been previously reported to be linked to a loss of IC^26–28^. Moreover, this could be explained by the loss of CD28 on T-cells, which has also been associated with increased circulating levels of inflammatory cytokines and biomarkers such as IL-6 and CRP^29^.

We also validated the ability of DNAm IC to capture physical and mental capabilities. We observed that individuals with high DNAm IC display improved pulmonary function (i.e. peak expiratory flow, forced vital capacity, forced expiratory volume). This observation is consistent with a recent study in the UK Biobank showing that lower levels of IC are associated with a significant increase in the increased risk of respiratory disease mortality^30^. Similarly, individuals with high DNAm IC exhibited improved bone mineral density (BMD) and engaged in more physical activity, which is known to contribute to BMD and overall physical resilience^31,32^. Interestingly, we found that individuals with higher DNAm IC consumed more fish and had higher levels in the blood of omega-3 fatty acids (DHA, docosahexaenoic acid; DPA, docosapentaenoic acid; EPA, eicosapentaenoic acid) abundant in marine sources. However, a 3-year clinical trial of omega-3 supplementation (800 mg DHA and 225 mg EPA per day) reported no improvement in overall IC^33^.

Despite our progress in predicting and understanding the molecular basis of IC, we discovered several areas for improvement. Although INSPIRE-T includes individuals up to 102 years old, IC rapidly declines at very old age (>90 years old), resulting in a very few individuals (2.9%) covering half of the potential decline in IC and rendering insufficient statistical power to make accurate predictions of very low IC. Also, despite identifying a strong association between IC and the immune system, it is still unclear if there are causal relationships underlying this association, especially considering that cytomegalovirus (CMV) infection might potentiate the expansion of CD28^-^ CD4^+^ T-cells^34,35^. We anticipate that future iterations of the INSPIRE-T cohort, incorporating longitudinal data and clinical parameters, will bridge these gaps by facilitating the identification of causal relationships between IC changes and immune function, as well as the molecular prediction of IC at all levels.

## Methods

### Intrinsic capacity score

To calculate the IC score, we used data from the INSPIRE-T cohort (version 1.0). Briefly, the INSPIRE-T cohort is an ongoing 10-year follow-up study investigating IC changes and biomarkers of aging and age-related diseases. Participants ranged from 20 to 102 years old and covered all levels of functional capacity. We calculated an overall IC score based on the following variables describing five health domains: Cognition: Mini-Mental State Examination (MMSE; score range 0-30, higher is better)^36^. Locomotion: Short Physical Performance Battery (SPPB; score range 0-12, higher is better)^37^. Psychology: Nine-item Patient Health Questionnaire for depression (PHQ-9; score range 0-27, higher is worse)^38^. Sensory: Visual acuity measured by the WHO simple eye chart (score range 0-3, higher is better) and hearing measured by the whisper test (score range 0-2, higher is better). Despite the WHO ICOPE Handbook recommending MMSE, SPPB, PHQ-9, WHO simple eye chart, and whisper test as tools to approximate domains of IC, there is no agreed-upon measure for vitality in the literature. According to the WHO, vitality can include factors related to energy, metabolism, neuromuscular function, and immune response. In this study, we used handgrip strength as a measure of vitality because it is a marker of physiological reserve and is associated with mortality across all ages^39,40^. From the 1,014 individuals in INSPIRE-T, 973 were assessed in all five domains of intrinsic capacity. Raw scores for each individual were re-scaled from 0 to 1, where higher is better (PHQ-9 scores were reversed to match the direction). Given the different score distributions, we z-transformed the values. The overall IC score was defined as the average of the z-transformed values across domains. We calculated the sensory domain score by averaging the visual and hearing scores. Finally, we performed min-max normalization on the overall IC score to provide an interpretable metric.

### DNA methylation data

We performed DNA methylation profiling (EPIC array) on 1,002 individuals in the INSPIRE-T cohort. We read the raw methylation data using the read.metharray.exp function from minfi^41^. We computed detection p-values to evaluate sample quality and filtered out samples with a high proportion of failed probes (average p > 0.01). We performed initial preprocessing using the preprocessRaw function and identified samples with outlier methylation profiles using QC plots. We removed CpG sites with poor detection p-values (p > 0.01) from any sample. We then converted the methylation dataset into a genomic ratio set and mapped it to the genome. We removed probes at CpG sites with SNPs and cross-reactive probes and excluded sex chromosome probes. We normalized beta values representing methylation levels using the BMIQ method^42^ implemented in the R package ChAMP^43^. We processed DNA methylation data from the FHS using the same method.

### Model generation

We used DNA methylation beta values and IC scores to build a predictive model for IC. To ensure robust predictions, we split the IC score into 20 bins of 0.05 and only considered bins with at least 10 individuals for the prediction. This criterion included individuals with an IC between 0.55 and 1. Therefore, we excluded samples with an IC below 0.55, which accounted for 2.9% of the total data. Using the glmnet package^44^, we performed a 10-fold cross-validated elastic net regression with alpha parameters ranging from 0.1 to 1, using mean absolute error (MAE) as the performance metric. We selected the optimal lambda in each model based on the minimum cross-validated error.

### Comparisons with epigenetic clocks

We compared our IC clock with established epigenetic clocks using IC predictions based on DNA methylation (DNAm IC) and epigenetic age estimates calculated with the methylclock package^45^, including the Horvath, Hannum, and Levine clocks. We computed residuals from linear models that adjusted for chronological age for each clock and DNAm IC (age-adjusted DNAm IC). We calculated Spearman’s correlation coefficients to evaluate the relationships between the IC clock and each epigenetic clock.

### Gene expression enrichment

Using the generated model, we predicted IC for individuals in the Offspring cohort of the FHS using DNA methylation data (obtained during the 8th exam) and investigated its association with gene expression changes, followed by enrichment analysis. We performed differential gene expression analysis using linear models to identify genes associated with age-adjusted DNAm IC. We filtered genes with an FDR < 0.05. We conducted enrichment analysis using the clusterProfiler package^46^ to identify biological processes associated with differentially expressed genes. We visualized enriched Gene Ontology (GO) terms using the CellPlot package (https://github.com/dieterich-lab/CellPlot).

### Cell count estimation

We estimated the proportions of CD8^+^ T-cells, CD4^+^ T-cells, natural killer cells, B cells, and granulocytes using Houseman’s estimation method via the meffilEstimateCellCountsFromBetas function implemented in the methylclock package. We integrated the age data and calculated correlations between cell counts and age using Spearman’s correlation coefficients. We also conducted correlation analyses between the age-adjusted cell counts and the age-adjusted DNA IC.

### Mortality analysis

We compared DNA methylation-based IC (DNAm IC) and traditional epigenetic clocks (Horvath, Hannum, and PhenoAge) for their ability to predict mortality using FHS data. We computed age-adjusted residuals for DNAm IC and each clock. We grouped the subjects into quintiles based on DNAm IC residuals, comparing the top and bottom 20%. We used Cox proportional hazards models^47^, adjusted for age, to calculate hazard ratios (HRs) and 95% confidence intervals (CIs) associated with mortality from all causes, cardiovascular diseases, congestive heart failure, and stroke/TIA. We plotted Kaplan-Meier survival curves to visualize survival differences between high and low IC groups, and calculated p-values using a log-rank test^48^.

### Association with health-related assessments and lifestyle

We analyzed the correlation between age-adjusted IC and various physiological and clinical measurements using data from the FHS, including clinic lab assays, inflammatory markers, Tau levels, cardiovascular risk factors, and dietary data. We calculated Spearman’s correlation coefficients (R) and p-values for the relationship between IC and each feature, applying false discovery rate (FDR) adjustments for multiple comparisons. Significant correlations (FDR < 0.05) were visualized using the ComplexHeatmap package^49^, and scatter plots were created for dietary factors (flavonoids, fatty acids, food frequency, and dietary guidelines adherence).

## Acknowledgements

We would like to thank the INSPIRE-T participants, and all the research and clinical staff who contributed to the study. The INSPIRE-T study was supported by grants from the Region Occitanie/Pyrénées-Méditerranée (Reference number: 1901175), the European Regional Development Fund (ERDF) (Project number: MP0022856), the Inspire Chairs of Excellence funded by: Alzheimer Prevention in Occitania and Catalonia (APOC), EDENIS, KORIAN, Pfizer, Pierre-Fabre, and the IHU HealthAge which received funding from the French National Research Agency (ANR) as part of the France 2030 program (reference number: ANR 23 IAHU 0011). The IHU HealthAge Open Science initiative was supported by the French National Research Agency (ANR) as part of the France 2030 program (reference number: ANR 23 IAHU 0011), and builds on the work conducted in the Data Sharing Alzheimer project.

